# Interregional human assembloids recapitulate fetal brain morphologies and enhance neuronal complexity

**DOI:** 10.1101/2025.10.24.684445

**Authors:** André Saraiva Leão Marcelo Antunes, Maria Carolina Pedro Athié, Ana Clara Caznok Silveira, Josué Renner, Elayne Vieira Dias, João Victor Ribeiro dos Santos, Branka Hrvoj-Mihic, Camila Canateli, João Meidanis, Alysson R Muotri, Alberto Antônio Rasia-Filho, Simoni Helena Avansini

## Abstract

Neuronal morphology governs how neurons connect, integrate, and process information, offering critical insights into the functional architecture of the brain. Characterizing the three-dimensional (3D) morphology of individual neurons is key not only for mapping circuit connectivity but also for understanding the cellular diversity that emerges during development. Neural organoids are valuable models of human brain development and disease, yet their morphological complexity remains poorly characterized despite advances in single-cell transcriptomics. Here, we use 3D confocal imaging and manual reconstruction of 735 neurons to analyze forebrain (dorsal and ventral) and thalamic (dorsal and ventral) organoids, as well as forebrain, thalamic, and corticothalamic assembloids. We find that organoids and assembloids exhibit distinct morphologies resembling fetal brain neurons, including immature pyramidal-like, double-bouquet, and bushy-like neurons. Interregional assembloids show greater neuronal morphological complexity than individual organoids, with more extensive dendritic branching, longer projections, and diverse soma shapes. Corticothalamic assembloids further display features of emerging connectivity. We observe dendritic spines with excitatory and inhibitory profiles and varicosities, indicative of maturing synaptic architecture. Together, our work makes an initial effort in describing the diversity of neuronal morphology in human neural organoids and assembloids. It further establishes structural phenotyping as a critical dimension for validating human neural models and underscores their value for modeling morphofunctional disorders.

## INTRODUCTION

Why does understanding the cellular morphology of a human brain model matter? Neuronal morphology governs how neurons connect, integrate, and process information, offering critical insights into the functional architecture of the brain.^1^ Characterizing the 3D morphology of individual neurons is therefore key not only for mapping circuit connectivity but also for understanding the cellular diversity that emerges during development. Variations in neuronal structure are known to contribute to functional heterogeneity across cell types, brain regions, developmental stages, and differential vulnerability to disease. ^2,3^

Three-dimensional neural organoids have emerged as powerful tools to model human brain development and neurological disorders. While single-cell transcriptomic studies have revealed substantial molecular diversity within organoids^4^, the morphological features of neurons - such as dendritic arborization, somatic morphology and synaptic specialization - have not been characterized.

To address this gap, we performed morphometric analyses of neurons in three-month-old, cleared organoids modeling distinct forebrain and thalamic regions and their interregional assembloids. We generated dorsal (Fd; pallium) and ventral (Fv; subpallium) forebrain organoids, which together mimic cortical development^5,6^, which were fused to generate forebrain assembloids (AF). We also generated dorsal (Td) and ventral (Tv) thalamic organoids, whose assembly recapitulates thalamic organization, giving rise to thalamic assembloids (AT).^7,8^ Dorsal structures are enriched in glutamatergic neurons, whereas ventral structures predominantly contain GABAergic neurons. Finally, combining AF and AT structures yielded corticothalamic assembloids (ACT), enabling investigation of emerging long-range connectivity (Figures 1A and S1). Together, these models allowed us to make an initial effort in describing the diversity of neuronal morphology in human neural organoids and assembloids.

**Figure 1.**
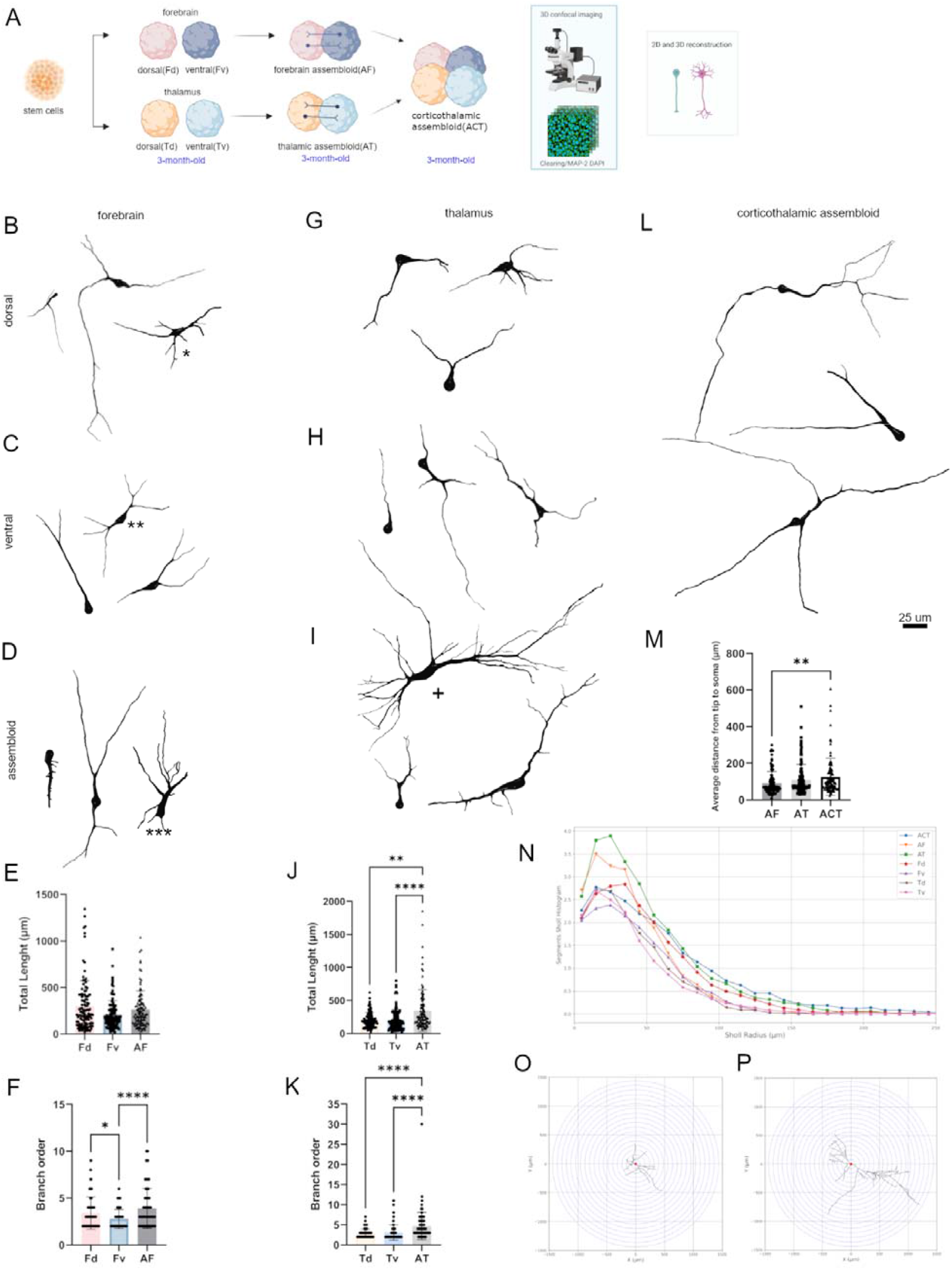
Morphological and morphometric characterization of region-specific human neural organoids and assembloids. (A) Schematic of the experimental workflow. Human pluripotent stem cell-derived organoids were patterned to resemble dorsal (Fd) and ventral (Fv) forebrain, and dorsal (Td) and ventral (Tv) thalamic regions. These were assembled to generate forebrain (AF), thalamic (AT), and corticothalamic (ACT) assembloids. All models were cultured for the same total duration (90 days); however, the timing of assembly varied: AF and AT were assembled at day 30 and maintained for 60 days, while ACT was generated at day 60 and cultured for the final 30 days. At the end of culture, samples were cleared, immunolabeled with MAP-2 and DAPI, and imaged using whole-mount 3D confocal microscopy with tiled z-stack acquisition. Manual reconstruction of individual neurons was performed using NeuTube. (B-D) Representative 3D reconstructions of neurons from dorsal forebrain organoids (B), ventral forebrain organoids (C), and forebrain assembloids (D). (E-F) Quantification of total neurite length (E) and branch order (F) in forebrain groups. (G-I) Representative reconstructions from dorsal thalamic (G), ventral thalamic (H), and thalamic assembloids (I). (J-K) Quantification of total neurite length (J) and branch order (K) in thalamic groups. (L) Representative reconstruction from a corticothalamic assembloid. (M) Quantification of average tip-to-soma distance across forebrain, thalamic, and corticothalamic assembloids. (N) Sholl analysis comparing arborization complexity across all groups, based on the mean number of segments at increasing distances from the soma. (O-P) Concentric segment analysis. (O) Representative Sholl profile from a ventral thalamic organoid-derived neuron. (P) Representative profile from a thalamic assembloid-derived neuron. A total of 735 neurons were reconstructed across groups: Fd (n = 111), Fv (n = 103), Td (n = 105), Tv (n = 107), AF (n = 102), AT (n = 105), ACT (n = 102). Data are presented as mean ± S.D. Statistical significance was determined using Kruskal-Wallis test followed by Dunn’s post hoc test: ^*^p < 0.05, ^**^p < 0.001, ^****^p < 0.0001. Putative neuronal subtypes are indicated in the representative reconstructions: immature pyramidal neuron (B), ^**^ double-bouquet cell (C), ^***^ basket cell-like neuron (D), bushy-like neuron (I). Scale bar: 25 μm.

## RESULTS

To illustrate the diversity of human neuronal morphology, we arranged a subset of neurons from region-specific organoids and assembloids along a continuum of increasing size and structural complexity, defined by neurite number, length, and branching patterns (Figures 1B-1D, 1G-1I, 1L, and S2).

Along this morphological continuum, MAP2-positive unipolar neurons typically extend a single process of variable length and thickness, ranging from unbranched to having four branches. Multipolar neurons with two processes show heterogeneous dendritic patterns. Those bearing three or more neurites exhibit diverse somatic morphologies - pyramidal, ovoid, fusiform - suggestive of distinct neuronal morphotypes. In some cases, the conical thinning arising from the cell body was suggestive of an axon hillock (Figure S2 asterisks). Soma shape and size are known to correlate with intrinsic electrophysiological properties and developmental stage, while dendritic architecture further influences somatic structure and the connectional and biophysical properties of the neuron.^9^

Across all morphological types, neuronal processes occasionally showed abrupt loss of fluorescent immunolabeling. At early stages, neurons typically displayed sparse arborization and multidirectional neurites with limited branching complexity - consistent with morphological features observed during *in vivo* development. Classification of embryonic neurons was challenging due to their dynamic maturation, initial simple morphology, and lack of laminar or nuclear landmarks. Neurons from forebrain and thalamic organoids exhibited distinct size and shape profiles along the continuum, resembling region-specific morphological subtypes reported for the cerebral cortex and thalamus in post-mortem studies.10,11 Assembly amplified neuronal morphological diversity beyond that seen in individual organoids. In both forebrain and thalamic assembloids, higher-order dendritic branching (second- and third-degree) appears even at intermediary stages of the continuum (Figures 1D, 1I, 1L, S2E-S2G and S3), whereas such features are rare or absent in multipolar neurons from individual organoids. Corticothalamic assembloids exhibit an additional level of complexity, with more extensive distal projections than those in either forebrain or thalamic assembloids alone.

### Assembly of Region-Specific Organoids Increases Neuronal Complexity and Reveals Subtype-Specific Neurons

Forebrain organoids contain immature uni- and multipolar neurons with ovoid or fusiform somata and variably branched neurites, some exhibiting immature pyramidal-like morphology. These neurons extend predominantly distal projections with a radial orientation. Ventral forebrain neurons, which are primarily GABAergic, are consistently smaller in both soma size and total neurite length compared to dorsal forebrain organoids (Figure 1E), a hallmark of developing inhibitory neurons.^12^ In forebrain assembloids, neuronal projections reach lengths comparable to those in dorsal forebrain organoids but exhibit more branched neurites, showing approximately a 1.4-fold increase in average branch order when compared to individual organoids (Figures 1E-1F). This pattern was further supported by Sholl analysis, which revealed a higher number of dendritic segments within 50 μm radius from the soma relative to forebrain organoids, indicating more branched dendrites in assembled forebrain models (Figures 1N-1P). These findings suggest that assembly enhances structural complexity of neurons. Multiple neuronal subtypes appear to be in development, including putative immature pyramidal neurons (Figure 1B, asterisk) and nonpyramidal subpopulations, such as presumed double-bouquet cells^12^ (Figure 1C, double asterisk) and basket cells^13^ (Figure 1D, triple asterisk).

Thalamic organoids predominantly contain neurons with polymorphic somata, typically oval or irregular, and neurites that follow tortuous trajectories. In ventral thalamic organoids, we observed putative interneurons with distinctive morphological features, including a small ovoid soma and short, branched processes bearing dendritic varicosities.^14^ Neurons from thalamic assembloids exhibited increased dendritic complexity compared to those from thalamic organoids, characterized by 1.7-fold increase in total length (Figures 1G-1I and 1J) and an approximately 1.6-fold increase in branch order (Figure 1K). Sholl analysis revealed a higher number of neuritic segments within the 150 μm radius from the soma compared to dorsal and ventral thalamic organoids (Figure 1N), along with a wider spread of the arbor, as indicated by the increased average distance from tip-to-soma (i.e. distal projections) in thalamic assembloids compared to individual thalamic organoids (Figure S3E). We also identified neurons with a bushy-like morphology (Figure 1I+), suggestive of a glutamatergic thalamic subtype previously described in human and rodent brains.^15^

Corticothalamic assembloids exhibited the greatest morphological diversity among all models, with neurons displaying ovoid, round, and elongated somata, as well as more extensive projections (Figures 1L-1M). These neurons possessed the longest neurites across all conditions, with a lower number of dendritic branches compared to those in forebrain and thalamic assembloids (Figure S3G-S3H). Despite this, their spread of dendritic arbor and total length remained comparable with other groups (Figures S3I and S3J), indicating that neurons in corticothalamic assembloids branch less frequently but extend their processes over longer distances. Sholl analysis revealed a higher number of segments beyond 100 μm from the soma, confirming their distinct dendritic organization characterized by more distal branching (Figure 1N). Notably, this structural complexity emerged over a shorter co-culture period of corticothalamic assembloids compared to forebrain and thalamic assembloids (see Methods). These observations are consistent with studies of human corticothalamic development, in which prefrontal neurons increase in length following the arrival of thalamic inputs.^14,16–18^

Altogether, our findings indicate that interregional assembly fosters greater structural complexity in organoid-derived neurons, as reflected by increased branching and extended projection length (Videos S1-S3). Sholl analysis further highlights this pattern: despite equal culture times, neurons from assembloids display significantly more elaborate arborization which extends farther along dendritic shafts when compared to individual organoids (Figures 1O-1P). Notably, while corticothalamic assembloids exhibited the highest overall morphological diversity, thalamic neurons showed the most substantial increase in neuritic complexity upon assembly, highlighting the differential responsiveness of distinct neuronal populations to interregional interaction. These results reinforce the role of regional assembly in promoting complex dendritic arborization and advancing neuronal maturation.

### Emergent Morphological Features in Assembloids: Dendritic Spines and Varicosities

Assembled complexity is reflected in the presence of dendritic filopodia and somatic and dendritic spines with diverse sizes and shapes - stubby and thin spine - appearing isolated or clustered nearby (Figure 2A). We observed increased dendritic spine density in assembloids, with spines frequently appearing in clusters - a pattern that may reflect enhanced synaptogenesis. Co-localized synaptic puncta confirmed the presence of both excitatory and inhibitory synapses on these developing neurons (Figure 2B). All models exhibited varicosities of varying sizes along neurites (Figures 2C-2E, Video S4), representing, to our knowledge, the first report of such structures in human neural organoids. Some enclosed varicosities contained vacuole-like structures^19^. These putative synaptic boutons were most abundant in ventral thalamic organoids and assembloids, consistent with the emergence of thalamic reticular components mediating local GABAergic inhibition.^20,21^ We also identified an uncommon morphological feature: a dendritic shaft that bifurcates and subsequently fuses several micrometers downstream, a configuration that may affect local signal propagation or branch-specific electrical resistance (Video S5).

**Figure 2.**
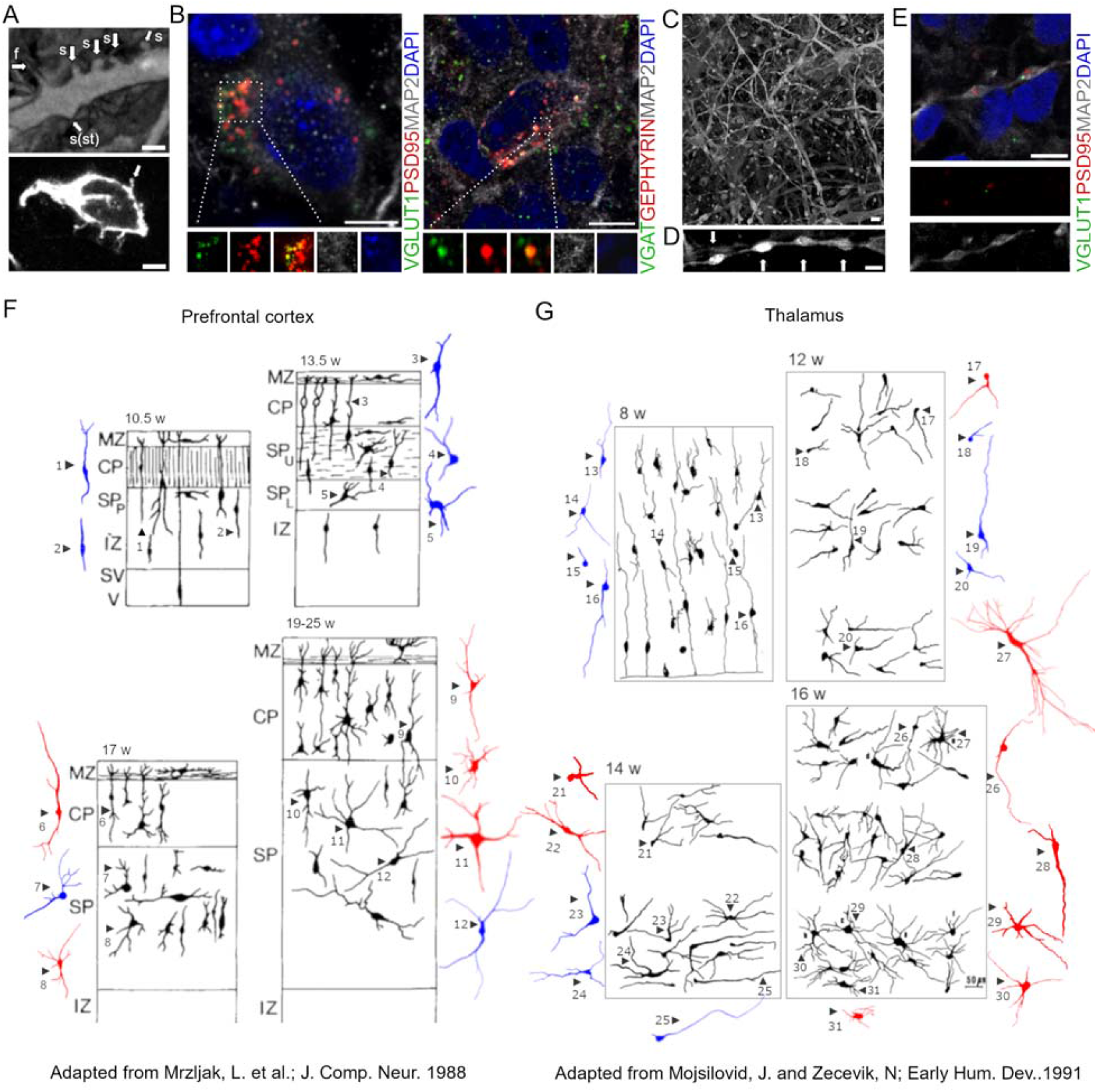
Dendritic spines and varicosities in neurons derived from assembloids and comparison of neuronal morphology of human fetal brain regions and corresponding organoids and assembloids. (A) Confocal microscopy images highlighting dendritic (upper image) and somatic (bottom image) spines in neurons from a ventral forebrain organoid (100 x), and forebrain assembloid (63 x), respectively. Note the presence of filopodia (f) and pleomorphic spines (s), including stubby (st) types, along with other spines with transitional morphologies. Scale bar: 5 μm (B) Representative co-localization of synaptic puncta (yellow) in a thalamic assembloid. Excitatory (left) and inhibitory (right) synapses are labeled with presynaptic markers VGLUT1 and VGAT (green), postsynaptic markers PSD-95 and Gephyrin (red), MAP-2 (gray), and DAPI (blue), respectively. Scale bar: 5 μm. (C) Two-dimensional maximum intensity projection depicting several varicosities along neuronal processes. Scale bar: 5 μm. (D) High-resolution image of a neurite segment highlighting the varicosity aspect, the inter-varicosity spacing, and their regularity (indicated by arrows). Scale bar: 5 μm. (E) Image showing putative excitatory synaptic markers in varicosities, labeled with presynaptic marker VGLUT1 (green), postsynaptic marker PSD-95 (red), MAP-2 (gray), and DAPI (blue). Scale bar: 100 μm. (F–G) Comparative schematics of neuronal development in the human fetal prefrontal cortex (F) and thalamus (G) across gestational stages (fetal images reprinted with permission), shown alongside corresponding features observed in forebrain and thalamic organoids and assembloids. Reconstructed neurons from the present study are overlaid in color to highlight morphological resemblance to fetal neurons (shown in black). Blue reconstructions represent neurons from region-specific organoids, while red reconstructions depict neurons from assembloids. Fd: dorsal forebrain organoids; Fv: ventral forebrai oorganoids; Td dorsal thalamic organoids; Tv: ventral thalamic organoids; AF: forebrain assembloids; AT: thalamic assembloids; and ACT: corticothalamic assembloids.

### Organoid-Derived Neuronal Morphologies Reflect Features of Human Fetal Brain

Organoid-derived neurons exhibited morphological features that resemble those described in the developing human cerebral cortex and thalamus based on fetal postmortem studies. Forebrain and thalamic organoids exhibited morphologies characteristic of early-stage development, corresponding to 10-13 and 8-12 gestational weeks, respectively.^11,16^ These included a predominance of simple unipolar and multipolar neurons with two processes. Consistent with prior transcriptomic analyses, organoids generated under similar conditions have been shown to match fetal brain tissue from the first trimester at the molecular level.^4^ In assembloids, neurons displayed features consistent with later developmental stages - approximately 17-25 gestational weeks for forebrain and 14-16 weeks for thalamus - including increased dendritic branching and more extended projections (Figures 2F-2G). These patterns align with previous descriptions of prenatal neuronal maturation.^11,16^ While cultured neurons develop in an environment distinct from that of the *in vivo* fetal brain, our findings support the capacity of regionally patterned organoids and assembloids to capture key structural hallmarks of neuronal development.

## DISCUSSION

Our single-neuron morphological analysis extends previous efforts to characterize neural organoids beyond molecular and physiological identity, revealing features reminiscent of human fetal brain development. Morphological maturation is enhanced in assembloids, which recapitulate key aspects of region-specific neuronal architecture.

Efforts to define neuronal diversity have underscored the importance of integrating molecular, physiological, and morphological properties. Connectivity represents a fourth, equally fundamental axis of neuronal identity that remains difficult to assess due to technical challenges.^22^ Our findings represent an initial step toward strengthening the morphological axis of organoid-based modeling. We show that regionally patterned organoids and assembloids generate neurons with diverse morphologies, including types consistent with immature pyramidal-like and bushy-like cells described in the fetal human brain. While transcriptomic and physiological profiling are well established, structural features remain undercharacterized.^4–8^ Our findings underscore the value of morphometric analysis as a complementary axis in organoid-based modeling, particularly for disorders involving altered neuronal morphology or connectivity.

### Limitations of the study

This study was performed using a single human pluripotent stem cell line, which may limit generalizability across genetic backgrounds. While our imaging pipeline enables detailed 3D reconstruction of neuronal morphology, higher-resolution and higher-throughput approaches will be essential to improve sampling depth and enable reliable quantification of fine structures. Additionally, neuronal classifications in this study were based solely on morphological criteria; integration with transcriptomic and electrophysiological modalities will be important for definitive cell type identification.

## Supporting information

Spreadsheet containing raw data of morphometric analyses

## RESOURCE AVAILABILITY

Requests for further information and resources should be directed to and will be fulfilled by the lead contact.

### Lead contact

Simoni Helena Avansini

### Materials availability

This study did not generate new unique reagents.

### Data and code availability

- Representative 3D confocal images (.tif) and reconstructed neuron files (.swc) reported in this paper have been deposited in the Zenodo repository under the DOI: 10.5281/zenodo.15756926 https://zenodo.org/records/15756926?preview=1&token=eyJhbGciOiJIUzUxMiJ9.eyJpZCI6ImQ1NmZhMjVmLWI2YzMtNGMyMi04YTNhLWMyZWYyMDNjZTE2ZiIsImRhdGEiOnt9LCJyYW5kb20iOiI4YWM3YTE4ZmQxNDYyMGUyNGU5Y2I2N2UxYTViNTg2YiJ9.0VU7FtuwTckLhWLsi7Q7ORUMJbTdHojRrGaBNxsICm5woI-PMRSSh88lNIb1pE6Bj1LwxrvWDkX41dF-ENp7Ig
- All original code has been deposited at GitHub and is publicly available at https://github.com/ana-caznok/Neuron_Morpho as of the date of publication.
- Any additional information required to reanalyze the data reported in this paper is available from the lead contact upon request.

## ACKNOWLEDGMENTS

We thank CNPEM (FNDCT-MCTI) for financial support and access to core research facilities. We are especially grateful to Stephanie Santos and Thayna Avelino (Biological Imaging Laboratory) for support with imaging, and to Pablo Silva and José Geraldo Pereira (Biological Data Team) for assistance with figure design and statistical analysis, respectively. This work was supported by an IBRO grant to S.H.A.; NIH grants R01MH100175, R01NS123642, R01AG078959, R01107788, R01AG084030, R01MH127077, R01DA056908, and R01MH123828, and DoD grant W81XWH2110306 to A.R.M.; CNPq/Brazil fellowship #314352/2020-1 to A.A.R.-F; and FAPESP grant 2024/01200-8 to J.M. We also thank Janaina Senna, Lila Pontes, Sandra Sánchez, and Srividya Ganapathy (UCSD) for their valuable support.

## AUTHOR CONTRIBUTIONS

A.S.L.M.A., M.C.P.A. and S.H.A. designed research; A.S.L.M.A., M.C.P. A., E.V.D., J.V.R.S., C.C. and S.H.A. performed research; A.S.L.M.A., M.C.P.A., A.C.C.S, J.R., E.V.D., J.V.R.S. B.H.M., C.C., J.M., A.A.R-F. and S.H.A. analyzed data; A.S.L.M.A., B.H.M., A.M., A.A.R-F. and S.H.A. wrote the paper.

## DECLARATION OF GENERATIVE AI AND AI-ASSISTED TECHNOLOGIES

During the preparation of this work, the authors used ChatGPT (OpenAI) for minor language editing. The authors reviewed and edited the content and takes full responsibility for the publication.

## SUPPLEMENTAL INFORMATION

**Document S1. Figures S1–S3, Table S1**

**Document S2. Spreadsheet containing raw data of morphometric analyses**

**Video S1: Representative 3D reconstruction of single neuron morphology derived from dorsal (left), ventral (middle) forebrain organoids and forebrain assembloid (right)**.

**VideoS2: Representative 3D reconstruction of single neuron morphology derived from dorsal (left), ventral (middle) thalamic organoids and thalamic assembloid (right)**.

**Video S3: Representative 3D reconstruction of single neuron morphology derived from assembloids. Neuron derived from forebrain assembloid (left); Neuron from thalamic assembloid (middle); Neuron from cortico-thalamic assembloid (right)**.

**Video S4: Representative 3D maximum intensity projection reconstruction of a cleared thalamic assembloid stained with MAP2, imaged with Leica LASX. The reconstruction reveals neurites containing varicosities of varying sizes along their length**.

**Video S5: Representative 3D maximum intensity projection of a cleared corticothalamic assembloid stained with MAP2, imaged with Leica LASX. The reconstruction shows a rare dendritic shaft that bifurcates and fuses several micrometers downstream, potentially affecting local signal propagation or branch-specific electrical resistance**.

## STAR★METHODS

### KEY RESOURCES TABLE

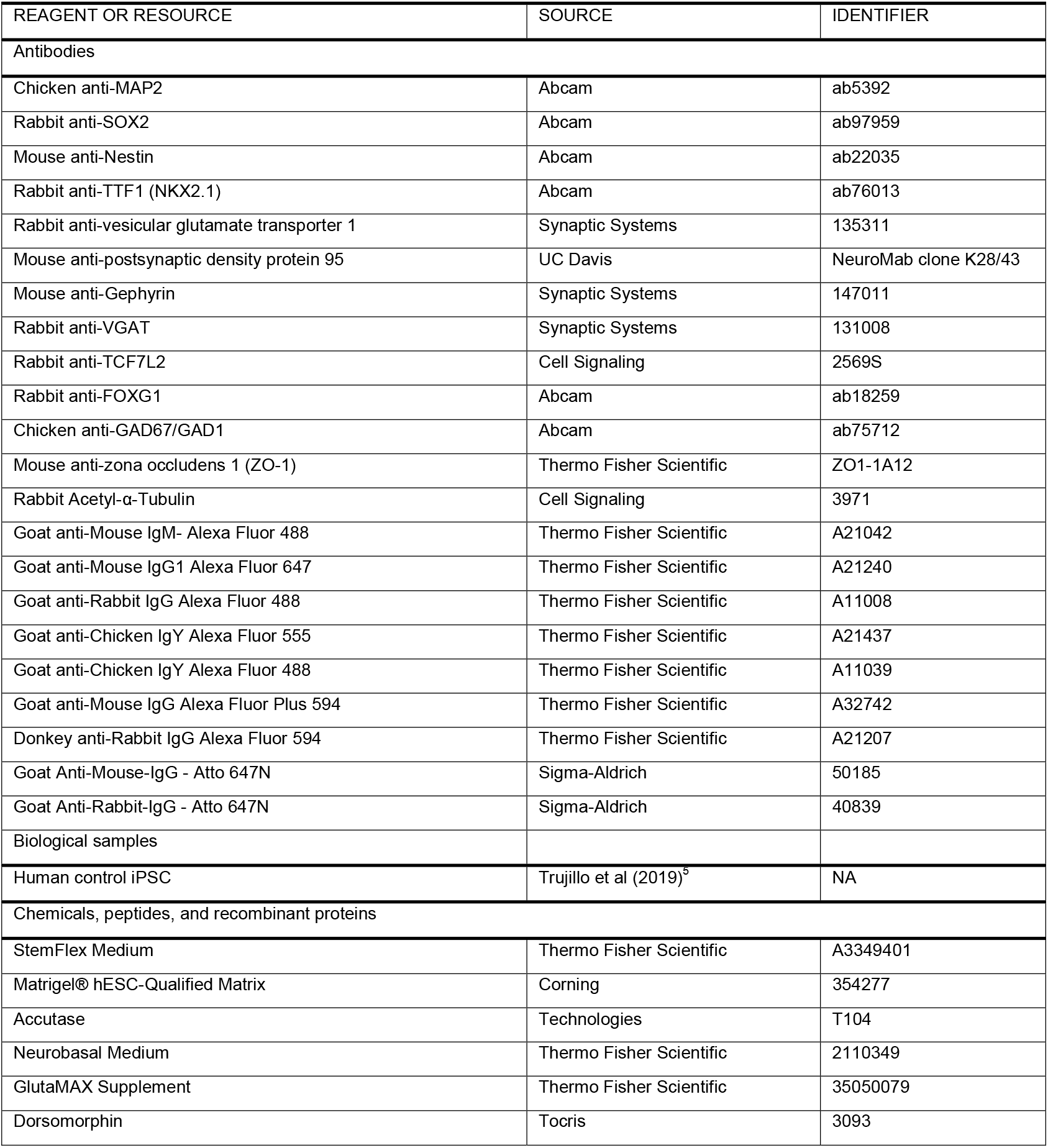
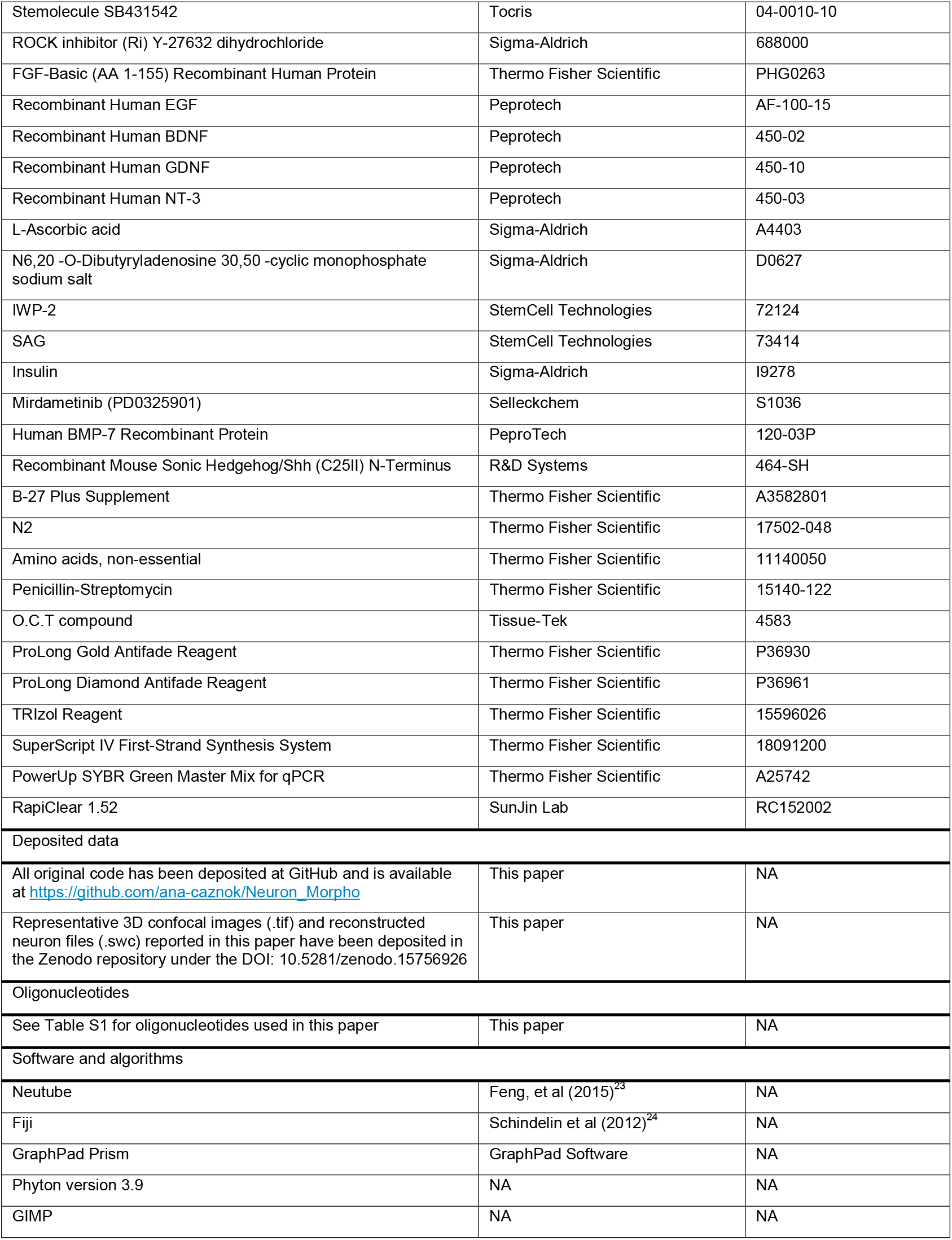

## EXPERIMENTAL MODEL AND STUDY PARTICIPANT DETAILS

### Human cell source

Human induced pluripotent stem cell (hiPSC) line from one male donor control individual used in this study were validated using methods described in previous studies^5^ (IRB Project #141223ZF). The iPSCs were expanded on Matrigel-coated dishes (Corning) in StemFlex medium (Thermo Fisher Scientific) and manually passaged. Pluripotency was confirmed by protein and gene expression analyses and by karyotyping with Infinium Global Screening Array BeadChip (Illumina Technologies). Cultures were routinely tested to ensure they remained free of Mycoplasma contamination.

## METHOD DETAILS

### Generation of region-specific human forebrain and thalamic organoids

To generate region-specific neural organoids, we followed established protocols for the dorsal forebrain,^5^ ventral forebrain,^6^ and dorsal and ventral thalamus^7,8^ with minor adaptations. The iPSCs were dissociated with Accutase (Thermo Fisher Scientific; T104) and aggregated in StemFlex medium (Thermo Fisher Scientific) supplemented with 10 µM SB431542, 1 µM dorsomorphin and 5 µM Rho kinase inhibitor (Y-27632) for 3 days. Approximately 4 × 10^6^ iPSCs were seeded into each well of a low-binding six-well plate and placed on a shaker inside a CO2 incubator at 95 rpm and 37°C. The formed spheres were subsequently transferred to neural induction medium, with specific details of each generation protocol are briefly described below.

Dorsal forebrain organoids were initiated by culturing spheres in neural induction medium containing 10 µM SB431542 (Tocris; 04-0010-10) and 1 µM Dorsomorphin (Tocris; 3093) for 7 days, followed by 14 days in expansion medium with 20 ng/mL FGF2 (7 days; Thermo Fisher Scientific; PHG0263) and then both 20 ng/mL FGF2 and 20 ng/mL EGF (7 days; Peprotech; AF-100-15). Neural maturation was induced using 10 ng/mL BDNF (Peprotech; 450-02), 10 ng/mL GDNF (Peprotech; 450-10), 10 ng/mL NT-3 (Peprotech; 450-03), 200 µM ascorbic acid (Sigma-Aldrich; A4403), and 1 mM dibutyryl-cAMP (Sigma-Aldrich; D0627) for 15 days, and organoids were then maintained in suspension under rotation in maintenance medium (5 ng/mL BDNF; 5 ng/mL NT-3;100 µM ascorbic acid).

Ventral forebrain organoids (subpallial) were initiated from day 6, when spheres were cultured in neural medium with 20 ng/mL EGF and 20 ng/mL FGF2 until day 24. Then, the neural medium was supplemented with 5 µM IWP-2 (days 4–24; StemCell Technologies; 72124), 100 nM SAG (days 12– 24; StemCell Technologies; 73414). From day 25 to 42, maturation was induced using 20 ng/mL BDNF and 20 ng/mL NT-3 and 200 µM ascorbic acid. From day 43, organoids were maintained in maintenance medium.

Dorsal and ventral thalamic organoids were initiated in neural induction medium containing 10 µM SB431542, 1 µM dorsomorphin and 4 µg/mL insulin (Sigma-Aldrich; I9278), for 3 days. Then, they were transferred to expansion medium with 20 ng/mL FGF2 (7 days) and then both 20 ng/mL FGF2 and 20 ng/mL EGF (7 days). From day 24 to 32, neural medium was supplemented using 30 ng/mL BMP7 (PeproTech;120-03P) and 1 µM PD325901(Selleckchem; S1036). From day 32 to 40, maturation was induced using 20 ng/mL BDNF and 20 ng/mL NT-3 and 200 µM ascorbic acid. From day 40, organoids were maintained in maintenance medium. For ventral thalamic organoids the neural induction medium was supplemented by adding 100 ng/mL SHH (from days 30–38).

Two independent batches of a single control iPSC clone were used for each region-specific organoid and assembloid. All models were cultured for 90 days, with assembloid formation initiated at approximately day 40 (forebrain and thalamic assembloids) or day 60 (corticothalamic assembloids). Regional identity was validated by immunofluorescence and gene expression profiling (please see Figure S1).

### Generation of assembloids

To generate forebrain assembloids, ventral and dorsal forebrain organoids were differentiated separately and then fused by placing them in close proximity in 96-well V-bottom plates (Corning) for 2 days in an incubator. The same procedure was used to form thalamic assembloids. For corticothalamic assembloids, preformed forebrain and thalamic assembloids were co-cultured in 96-well V-bottom plates (Corning) for 3 days to allow integration. Following assembly, all assembloids were transferred to ultra-low attachment six-well plates and maintained with medium changes every 4 days. The assembloids were cultured in neural maintenance medium until 90 days. The timing of assembly varied: forebrain and thalamic assembloids were generated around day 30 and cultured for 60 days, while corticothalamic assembloids were assembled on day 60 and maintained for an additional 30 days (Figure 1A).

### Immunofluorescence Staining

#### Whole-Mount Clearing and Immunostaining

Region-specific organoids and assembloids were fixed in 4% paraformaldehyde (PFA) overnight at 4 °C and stored in 1x PBS containing 0.02% sodium azide. After PBS wash, samples were permeabilized with 2% Triton X-100 in PBS for three days at 37 °C on an orbital shaker. Blocking was performed for two days at 4 °C using a solution of 10% normal goat serum, 1% Triton X-100, 2.5% DMSO, and 0.05% sodium azide in PBS. Organoids and assembloids were incubated with chicken anti-MAP2 antibody (Abcam, ab5392; 1:500) for five days at 4 °C under constant agitation. Goat anti-Chicken Alexa Fluor 555 (Thermo Fisher Scientific; A21437; 1:500) was applied for three days at 4 °C, followed by washes with PBS containing 3% NaCl and 0.2% Triton X-100. DAPI (1 µg/mL) was used for nuclear counterstaining (overnight at 4 °C). Samples were cleared using RapiClear 1.52 (SUNJing Lab, RC152002) at room temperature overnight) and mounted using iSpacer microchambers filled with fresh RapiClear, according to the manufacturer’s instructions.

### Cryosection-Based Immunostaining

Organoids and assembloids were fixed in 4% PFA overnight at 4 °C, cryoprotected in 30% sucrose for a minimum of two days, embedded in Tissue-Tek (Leica Microsystems), and sectioned at 20 µm on a Leica VT1000S cryostat. Sections were air-dried for 10 min, permeabilized in 1x PBS containing 1% Triton X-100 for 2 min and blocked with 1× PBS containing 0.1% Triton X-100 and 3% BSA for 3 h at room temperature. Slides were incubated overnight at 4 °C with the following primary antibodies: rabbit anti-SOX2 (Abcam; ab97959; 1:1000); mouse anti-NESTIN (Abcam; ab22035; 1:200); mouse anti-zona occludens 1 (ZO-1;Thermo Fisher Scientific; ZO1-1A12; 1:500); rabbit acetyl-α-tubulin (Cell Signaling; 3971; 1:200); Rabbit anti-TTF1 (NKX2.1; Abcam; ab76013; 1:200); rabbit anti-vesicular glutamate transporter 1 (VGLUT1; Synaptic Systems; 135311; 1:100); mouse anti-postsynaptic density protein 95 (PSD-95; NeuroMab, UC Davis, 1:100); mouse anti-Gephyrin (Synaptic Systems; 147011; 1:100); rabbit anti-VGAT (Synaptic Systems;131008; 1:100); rabbit anti-TCF7L2 (Cell Signaling; 2569S; 1:100); rabbit anti-FOXG1(Abcam: ab18259; 1:200) chicken anti-GAD1(Abcam; ab75712; 1:50). Following three washes in 1x PBS (5 min each), slides were incubated with secondary antibodies for 3 h at room temperature. Secondary antibodies used in this study were: Goat anti-Mouse IgM-Alexa Fluor 488 (Thermo Fisher Scientific; A21042; 1:500); Goat anti-Mouse IgG1 Alexa Fluor 647 (Thermo Fisher Scientific; A21240; 1:500); Goat anti-Rabbit IgG Alexa Fluor 488 (Thermo Fisher Scientific; A11008; 1:500); Goat anti-Chicken IgY Alexa Fluor 555 (Thermo Fisher Scientific; A21437; 1:500); Goat anti-Chicken IgY Alexa Fluor 488 (Thermo Fisher Scientific; A11039; 1:100); Goat anti-Mouse IgG Alexa Fluor Plus 594 (Thermo Fisher Scientific; A32742; 1:100); Donkey anti-Rabbit IgG Alexa Fluor 594 (Thermo Fisher Scientific; A21207; 1:100); Goat Anti-Mouse-IgG - Atto 647N (Sigma-Aldrich; 50185; 1:100); Goat Anti-Rabbit-IgG - Atto 647N (Sigma-Aldrich; 40839; 1:100). After further PBS washes, DAPI (1 µg/mL) was applied for 5 min, and sections were mounted using ProLong Gold or Diamond (Thermo Fisher Scientific). Confocal images were acquired using a DMi8 S Inverted Confocal Microscope (Leica).

### Real-time quantitative PCR (qPCR)

Total RNA was isolated from the whole organoids/assembloids using TRIzol (Thermo Fisher Scientific) following the manufacturer’s recommendations. Subsequently, 1.0 µg of total RNA was reverse transcribed into cDNA using the SuperScript® IV Reverse Transcriptase with DNase treatment included in the reaction (Thermo Fisher Scientific). For the quantification of gene expression, qPCR was carried out using QuantStudio 5 Real-Time PCR Systems (Thermo Fisher Scientific) and Power SYBR™ Green PCR Master Mix (Thermo Fisher Scientific), following the manufacturer’s instructions. Following PCR amplifications, a melting curve was analyzed for each gene, and a single melting temperature peak was observed. The PCR conditions were: 50 °C for 2 min, 95 °C for 2 min, followed by 40 two-step cycles at 95 °C for 15 s and 60 °C for 1 min. A list of primers used is presented in Table S1. PCR reactions were performed in triplicate. Relative expression was calculated using the 2^−ΔΔCt^ method, after normalization to endogenous control.

### 3D Imaging and Neuronal Reconstruction

Human neural organoids and assembloids were immunostained with MAP2 to label neuronal processes and enable morphological reconstruction. Three-dimensional imaging was performed using a DMi8 S Inverted Confocal Microscope (Leica) equipped with a 63x oil-immersion objective (HC PL Apo CS2 63x/1.40), with high-resolution tiled z-stack acquisition across optically cleared samples. Manual 3D reconstructions of individual neurons were performed using NeuTube (Center for Functional Connectomics, South Korea), an open-source software for morphological tracing, following previously established protocols.^25^ All individuals involved in reconstruction underwent calibration and cross-validation to ensure consistent criteria for tracing quality, branch resolution, and segmentation of neuronal processes. Neurons were randomly selected for reconstruction based on the clear visibility of a distinguishable soma and favorable positioning within the z-stack, such that the full or near-complete extent of the neurites could be captured in all directions. Each reconstruction included the soma and all visibly traceable neurites - defined here as cellular processes emerging from the soma whose dendritic or axonal identity could not be determined with certainty. Morphological classification was based on cell body shape and the number of primary processes: Unipolar neurons had a single neurite emerging from the soma. Multipolar neurons had two or more primary neurites. Cells with two primary processes - where one appeared to extend as a presumptive axon while the other retained MAP2 positivity - were designated as bipolar, following classical definitions by Ramón y Cajal (1909–1911).^19^ In cases where process identity could not be confidently assigned (e.g., due to short, straight projections without a visible hillock or tapering), the term “neurite” was retained. A total of 735 neurons were reconstructed across all experimental conditions, with quantification of soma morphology, total neurite length, branch order, and spatial distribution of branches.

### Morphometric and Statistical Analysis

Morphometric parameters were extracted using a novel graph-based analysis pipeline implemented in a custom Python code. The code, freely available at https://github.com/ana-caznok/Neuron_Morpho, takes as input.swc files containing a traced image with multiple neurons. It automatically separates the individual neuronal arbors, performs quantitative morphometric analysis, and generates plots of the extracted features. The code computes a series of biologically meaningful metrics. (1) Total neurite length was calculated by summing the lengths of all neurite segments for each neuron, providing a measure of the total length of all neurites per neuron. This metric can yield similar values for neurons with different structural profiles. For example, a small neuron with many short branches may have a total length comparable to that of a neuron with long unbranched neurites. (2) Number of branches and tips was assessed by identifying bifurcation points and terminal endpoints, respectively. To capture arbor organization, (3) branch order was computed by counting the number of bifurcations encountered along the shortest path from each node to the soma. This path was traced using a Depth-First Search (DFS) algorithm, which traverses the graph from the soma towards distal tips preserving its structure. This approach follows the centrifugal branch ordering convention used in postmortem studies,^26^ where primary branches emerging directly from the soma are designated as first-order branches, and each subsequent bifurcation increases the branch order incrementally (e.g., second-order, third-order, and so on).

Additionally, (4) average distance from branches and tips to the soma was calculated by applying a Breadth-First Search (BFS) algorithm to determine the shortest path from each branch point and terminal tip back to the soma. These distances were then averaged, providing a spatial measure of how far a neuron extends and at what distance from the soma branching typically occurs. Finally, (5) Sholl analysis is used to characterize the neuron arbor distribution as a function of radial distance from the soma.

Instead of counting intersections with individual circles, our approach quantified the number of neurite segments, branches, and tips located within successive radial shells defined between pairs of concentric spheres spaced 10 micrometers apart. Segments are defined as the region between two branching points or between a branching point and a tip. Quantification of neuronal morphology started by identifying the soma. In order to ensure consistency and biological accuracy across samples, the soma was defined as the node with the largest radius. Once identified, this node was designated as the graph root, and the neuronal structure was reoriented accordingly, ensuring that all path-based calculations reflected true spatial distances from the soma.

## QUANTIFICATION AND STATISTICAL ANALYSIS

Data are presented as mean ± standard deviation (S.D.). For morphometric parameters derived from reconstructed neurons within each region-specific organoid and assembloid, statistical significance was assessed using the Kruskal–Wallis test followed by Dunn’s post hoc test. Gene expression data were analyzed using one-way ANOVA with Tukey’s post hoc test. All analyses were performed using GraphPad Prism 9 (GraphPad Software Inc.). Statistical significance was defined as follows: ^*^p < 0.05, ^**^p < 0.01, ^***^p < 0.001 or ^****^p < 0.0001.

## REFERENCES

1. Luo, L. (2021). Architectures of neuronal circuits. Preprint, 10.1126/science.abg7285 10.1126/science.abg7285.

2. Liu, Y., Jiang, S., Li, Y., Zhao, S., Yun, Z., Zhao, Z.H., Zhang, L., Wang, G., Chen, X., Manubens-Gil, L., et al. (2024). Neuronal diversity and stereotypy at multiple scales through whole brain morphometry. Nature Communications 15. 10.1038/s41467-024-54745-6.

3. Peng, H., Xie, P., Liu, L., Kuang, X., Wang, Y., Qu, L., Gong, H., Jiang, S., Li, A., Ruan, Z., et al. (2021). Morphological diversity of single neurons in molecularly defined cell types. Nature 598, 174–181. 10.1038/s41586-021-03941-1.

4. Camp, J.G., Badsha, F., Florio, M., Kanton, S., Gerber, T., Wilsch-Bräuninger, M., Lewitus, E., Sykes, A., Hevers, W., Lancaster, M., et al. (2015). Human cerebral organoids recapitulate gene expression programs of fetal neocortex development. Proc Natl Acad Sci U S A 112. 10.1073/pnas.1520760112.

5. Trujillo, C.A., Gao, R., Negraes, P.D., Gu, J., Buchanan, J., Preissl, S., Wang, A., Wu, W., Haddad, G.G., Chaim, I.A., et al. (2019). Complex Oscillatory Waves Emerging from Cortical Organoids Model Early Human Brain Network Development. Cell Stem Cell 25. 10.1016/j.stem.2019.08.002.

6. Birey, F., Andersen, J., Makinson, C.D., Islam, S., Wei, W., Huber, N., Fan, H.C., Metzler, K.R.C., Panagiotakos, G., Thom, N., et al. (2017). Assembly of functionally integrated human forebrain spheroids. Nature 545. 10.1038/nature22330.

7. Xiang, Y., Tanaka, Y., Cakir, B., Patterson, B., Kim, K.Y., Sun, P., Kang, Y.J., Zhong, M., Liu, X., Patra, P., et al. (2019). hESC-Derived Thalamic Organoids Form Reciprocal Projections When Fused with Cortical Organoids. Cell Stem Cell 24. 10.1016/j.stem.2018.12.015.

8. Kiral, F.R., Cakir, B., Tanaka, Y., Kim, J., Yang, W.S., Wehbe, F., Kang, Y.J., Zhong, M., Sancer, G., Lee, S.H., et al. (2023). Generation of ventralized human thalamic organoids with thalamic reticular nucleus. Cell Stem Cell 30. 10.1016/j.stem.2023.03.007.

9. Udvary, D., Harth, P., Macke, J.H., Hege, H.C., de Kock, C.P.J., Sakmann, B., and Oberlaender, M. (2022). The impact of neuron morphology on cortical network architecture. Cell Rep 39. 10.1016/j.celrep.2022.110677.

10. Marín-Padilla, M. (2011). Human Motor Cortex Excitatory–Inhibitory Neuronal Systems: Development and Cytoarchitecture. In The Human Brain 10.1007/978-3-642-14724-1_6.

11. Mojsilovid, J., and Zecevik, N. (1991). Early development of the human thalamus: Golgi and Nissl study.

12. Somogyi, P., and Cowey, A. (1984). Double Bouquet Cells. In Cerebral Cortex Functional Properties of Cortical Cells, E. G. Jones and A. Peters, eds. (Springer New York, NY), pp. 337–360.

13. Jones, E.G., and Hendry, S.H.C. (1984). Basket Cells. In Cerebral Cortex Functional Properties of Cortical Cells, E. G. Jones and A. Peters, eds. (Springer New York, NY), pp. 309–336.

14. Bandeira, S., and Molnár, Z. (2022). Development of the Thalamocortical Systems. In The Thalamus, M. M. Halassa, ed. (Cambridge University Press), pp. 139–162. 10.1017/9781108674287.

15. Roy, D.S., Zhang, Y., Halassa, M.M., and Feng, G. (2022). Thalamic subnetworks as units of function. Preprint, 10.1038/s41593-021-00996-1 10.1038/s41593-021-00996-1.

16. Mrzljak, L., Uylings, H.B.M., Kostovic, I., and van Eden, C.G. (1988). Prenatal development of neurons in the human prefrontal cortex: I. A qualitative Golgi study. Journal of Comparative Neurology 271. 10.1002/cne.902710306.

17. Mrzljak, L., Uylings, H.B.M., Van Eden, G.G., and Judáš, M. (1991). Chapter 9 Neuronal development in human prefrontal cortex in prenatal and postnatal stages. Prog Brain Res 85. 10.1016/S0079-6123(08)62681-3.

18. Antón-Bolaños, N., Espinosa, A., and López-Bendito, G. (2018). Developmental interactions between thalamus and cortex: a true love reciprocal story. Preprint, 10.1016/j.conb.2018.04.018 10.1016/j.conb.2018.04.018.

19. Ramón y Cajal, S. (1911). Histologie du systeme nerveux de l’homme et des vertebres. Paris Maloine 2.

20. Pinault, D. (2023). The Thalamic Reticular Nucleus: Anatomo-Functional Mechanisms and Concept. In The Cerebral Cortex and Thalamus, W. M. Usrey and S. M. Sherman, eds. (Oxford University PressNew York), pp. 163–176. 10.1093/med/9780197676158.001.0001.

21. Südhof, T.C. (2021). The cell biology of synapse formation. Preprint, 10.1083/jcb.202103052 10.1083/jcb.202103052.

22. Zeng, H., and Sanes, J.R. (2017). Neuronal cell-type classification: Challenges, opportunities and the path forward. Preprint, 10.1038/nrn.2017.85 10.1038/nrn.2017.85.

23. Feng, L., Zhao, T., and Kim, J. (2015). Neutube 1.0: A new design for efficient neuron reconstruction software based on the swc format. eNeuro 2. 10.1523/ENEURO.0049-14.2014.

24. Schindelin, J., Arganda-Carreras, I., Frise, E., Kaynig, V., Longair, M., Pietzsch, T., Preibisch, S., Rueden, C., Saalfeld, S., Schmid, B., et al. (2012). Fiji: An open-source platform for biological-image analysis. Preprint, 10.1038/nmeth.2019 10.1038/nmeth.2019.

25. Guerra, K.T.K., Renner, J., Vásquez, C.E., and Rasia-Filho, A.A. (2023). Human cortical amygdala dendrites and spines morphology under open-source three-dimensional reconstruction procedures. Journal of Comparative Neurology 531. 10.1002/cne.25430.

26. C. J. Cupp, E. Uemura,. (1980) Age-related changes in prefrontal cortex of Macaca mulatta: Quantitative analysis of dendritic branching patterns. Exp Neurol 69. 10.1016/0014-4886(80)90150-8

