## Supplemental Data 1 for "Interregional human assembloids recapitulate fetal brain morphologies and enhance neuronal complexity"

**Document S1.**

**Figure S1**

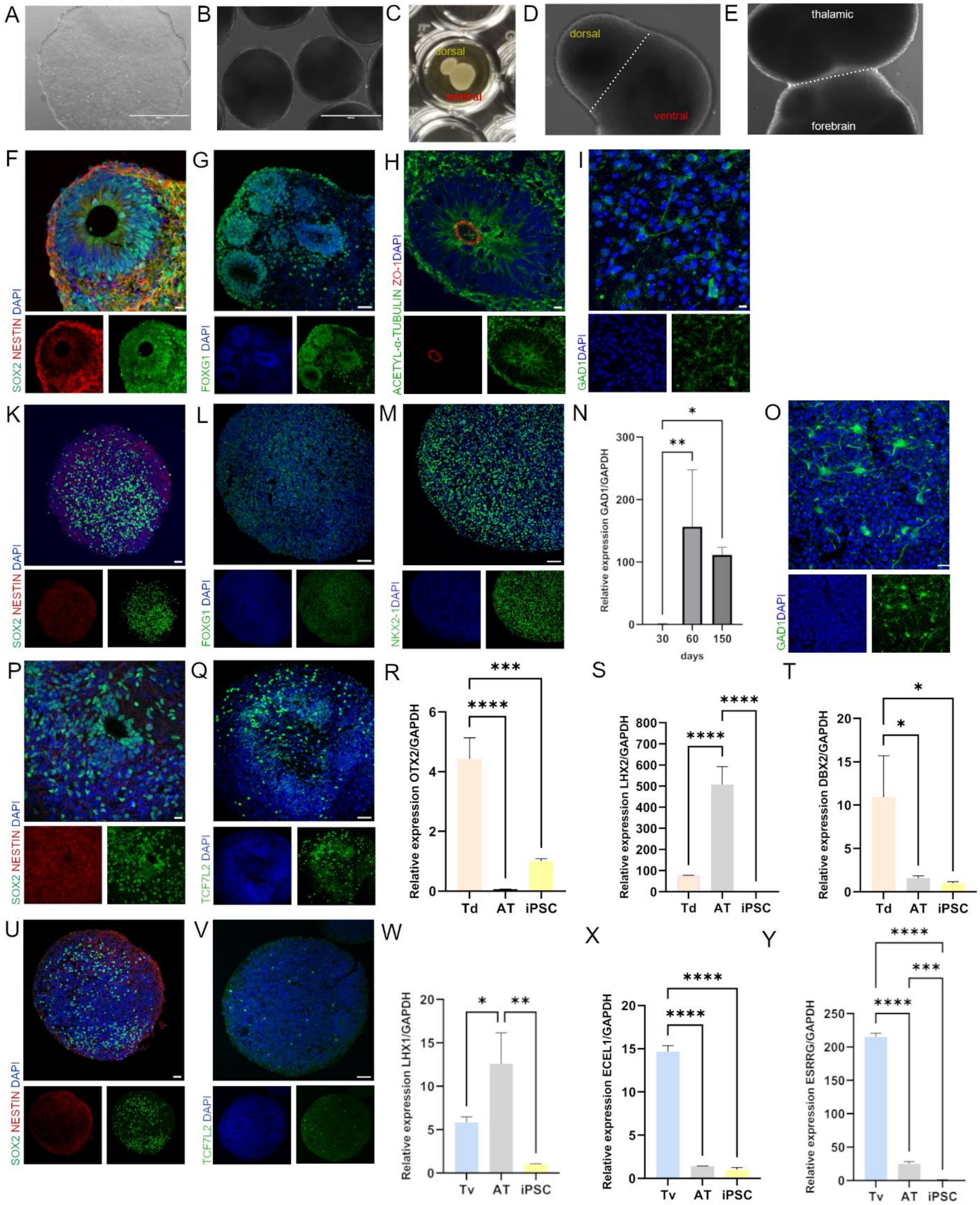


**Figure S1. Molecular characterization of human forebrain and thalamic neural organoids and assembloids.**

(A-B) Representative bright-field images of a control-derived human induced pluripotent stem cell (hiPSC) colony (A) and a 30-day-old dorsal forebrain organoid (B).

(C) Fusion of dorsal and ventral forebrain organoids to generate a forebrain assembloid.

(D-E) Bright-field images of a dorsal-ventral thalamic assembloid (D) and a four-part cortico-thalamic assembloid (E).

(F-I) Characterization of dorsal forebrain organoids: immunostaining for SOX2 (green) and Nestin (red) in neural progenitor cells of 30-day-old organoids (F); FOXG1 (green), a forebrain-enriched marker (G); ZO-1 (red) and Acetyl-α Tubulin (green) in 30-day-old organoids (H);  and GAD1 (green) in 180-day-old forebrain assembloids (I).

(K–O) Characterization of ventral forebrain organoids: SOX2 (green) and Nestin (red) (K); FOXG1 (green) (L); the medial ganglionic eminence marker NKX2-1 (green) (M); Quantification of *GAD1* expression at days 30, 60, and 150 (N); and GAD1 (green) in 300-day-old organoids (O).

(P-T) Characterization of dorsal thalamic organoids: SOX2 (green) and Nestin (red) in 52-day-old organoids (P); TCF7L2 (green) expression (Q); qPCR analysis showing enrichment of dorsal thalamic identity markers *OTX2* (R), *LHX2* (S), and *DBX2* (T) compared to iPSCs and 30-day-old thalamic assembloids.

(U-Y) Characterization of ventral thalamic organoids: SOX2 (green) and Nestin (red) in 52-day-old organoids (U); TCF7L2 (green) expression in 52-day-old organoids (V); qPCR showing expression of ventral markers *LHX1* (W), and thalamic ventral nuclear markers *ECEL1* (X) and *ESRRG* (Y), relative to iPSCs and 30-day-old thalamic assembloids.

Nuclei were counterstained with DAPI (blue). Gene expression data are shown as mean ± S.D. from three technical replicates. One-way ANOVA with Tukey's post hoc test was used to assess statistical significance; values were normalized to *GAPDH*. $^{*}p < 0.05$, $^{**}p < 0.001$, $^{****}p < 0.0001$. Abbreviations: Fd: dorsal forebrain organoid; Fv: ventral forebrain organoid; Td: dorsal thalamic organoid; Tv: ventral thalamic organoid; AF: forebrain assembloid; AT: thalamic assembloid; ACT: corticothalamic assembloid; iPSC, induced pluripotent stem cell. Scale bars: (A) 1000 µm; (B) 400 µm; (F-I), (K-M,O), (P-Q), (U-V): 50 µm.

**Figure S2**

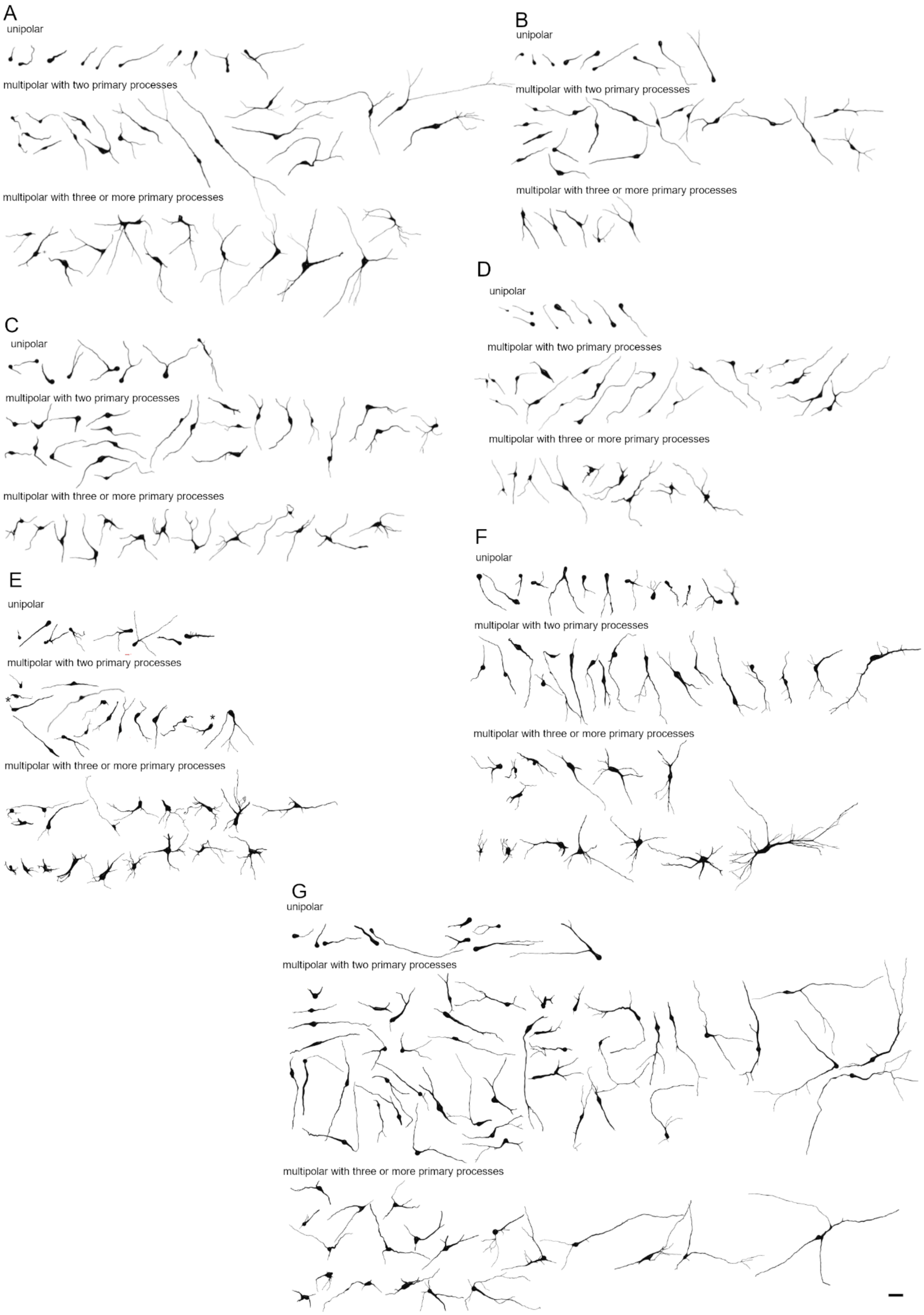
A
unipolar
multipolar with two primary processes
multipolar with three or more primary processes
B
unipolar
multipolar with two primary processes
multipolar with three or more primary processes
C
unipolar
multipolar with two primary processes
multipolar with three or more primary processes
D
unipolar
multipolar with two primary processes
multipolar with three or more primary processes
E
unipolar
multipolar with two primary processes
*
*
multipolar with three or more primary processes
F
unipolar
multipolar with two primary processes
multipolar with three or more primary processes
G
unipolar
multipolar with two primary processes
multipolar with three or more primary processes

**Figure S2. The morphological continuum of neuronal structural complexity in region-specific human neural organoids and assembloids.** (A–G) Representative 3D-reconstructed neurons from human region-specific neural organoids and assembloids arranged from unipolar to multipolar neurons with two or more processes from dorsal (Fd; A) and ventral (Fv; B) forebrain; dorsal (Td; C) and ventral (Tv; D) thalamic organoids; forebrain assembloid (AF; E); thalamic assembloid (AT; F); and corticothalamic assembloid (ACT; G). * indicate putative axon hillock. Note the variability in cell body shape shape and size, number of primary dendrites and branching pattern, and arbor architecture. Redundant morphologies were excluded for clarity. Scale bar: 25 μm.

**Figure S3**

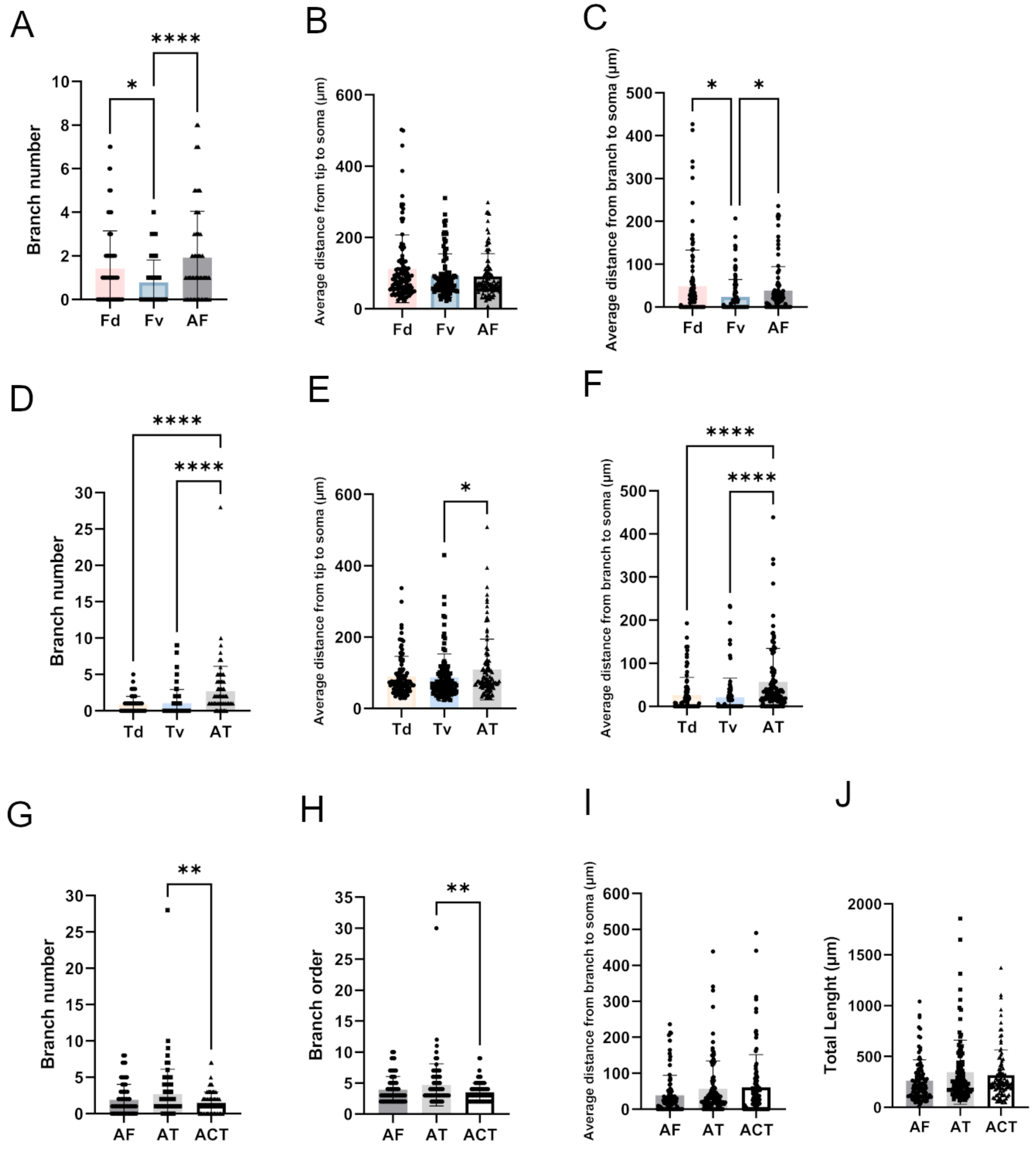


**Figure S3. Morphometric characterization of human region-specific neural organoids and assembloids.**

(A-C) Morphometric analysis comparing dorsal and ventral forebrain organoids and forebrain assembloids: branch number (A); average distance from tip to soma (B); average distance from branch to soma (C).

(D-F) Morphometric analysis comparing dorsal and ventral thalamic organoids and thalamic assembloids: branch number (D); average distance from tip to soma (E); average distance from branch to soma (F).

(G-J) Morphometric analysis comparing assembloids entities: branch number (G); branch order (H); average distance from branch to soma (I) and total neurite length (μm) (J).

A total of 735 neurons were reconstructed across groups: Fd (n = 111), Fv (n = 103), Td (n = 105), Tv (n = 107), AF (n = 102), AT (n = 105), ACT (n = 102). Data are presented as mean ± S.D. Statistical significance was determined using Kruskal-Wallis test followed by Dunn's post hoc test: *$p < 0.05$, **$p < 0.001$, ****$p < 0.0001$. Fd: dorsal forebrain organoids; Fv: ventral forebrain organoids; Td: dorsal thalamic organoids; Tv: ventral thalamic organoids; AF: forebrain assembloids; AT: thalamic assembloids; and ACT: corticothalamic assembloids.

**Table S1. Primer sequences used for gene expression analysis in region-specific organoids and assembloids.**

| Target | Sequences 5'-3' |
|---|---|
| *GAPDH* | Fwd: CCAGAAGACTGTGGATGGCC Rev: GGGATGATGTTCTGGAGAGCC |
| *GAD1* | Fwd: AGAGGGAACTAGCGAGAACG Rev: CAGAAACAGGCTCGGCTC |
| *OTX2* | Fwd: CCTCACTCGCCACATCTACTTT Rev: TGGAGCAGTGGAACTTACAGC |
| *LHX2* | Fwd: GGGTCCTCCAGGTCTGGTT Rev: CCGTGTTTTCCTGCCGTAAG |
| *DBX1* | Fwd: AGTACATCAGCAAGCCCGAC Rev: ACCAGATTTTCACCTGCGAGT |
| *LHX1* | Fwd: CCTCAACATGCGCGTCATTC Rev: CTCAGCTGCTTCATCCTCCG |
| *ECEL1* | Fwd: AGAACCAGATGGTGTTCCCC Rev: ATGCCCCCGTAGTTGAGAGA |
| *ESRRG* | Fwd: TCCCTTCTGAGAGAGCTGACA Rev: TGCGACAGTGTTAGCGTAGG |
